# Robust Serum Proteomic Signatures of APOE2

**DOI:** 10.1101/2025.05.24.655950

**Authors:** Paola Sebastiani, Eric Reed, Kevin B Chandler, Prisma Lopez, Hannah Lords, Harold Bae, Catherine E Costello, Matthew Au, Lingyi Lynn Deng, Mengze Li, Qingyan Xiang, Heeju Noh, Lance Pflieger, Cory Funk, Noa Rappaport, Marianne Nygaard, Meghan I Short, Michael Brent, Stefano Monti, Stacy L Andersen, Thomas T Perls

## Abstract

We previously identified a signature of 16 serum proteins that highlighted a role of the e2 allele of APOE in lipid regulation via apolipoprotein B (APOB) and apolipoprotein E (APOE), and in inflammation. The serum proteins were profiled using the aptamer-based Somalogic technology. Here, we validate and expand the serum protein signature of APOE using a combination of mass-spectrometry, ELISA, Luminex, antibody-based Olink proteomics, and blood transcriptomics. We replicate the association between APOB and the e2 allele of APOE, we correct the pattern of association between APOE genotypes and serum level of APOE, and we detect new associations between APOE genotypes and the complex of apolipoproteins APOC1, APOC4, APOC2, APOC3, APOE, APOF and APOL1. In addition, we discover 13 new proteins that correlate with APOE genotypes. This extended signature includes granule proteins CAMP, CTSG, DEFA3, and MPO secreted from neutrophils and points to olfactomedin 4 (OLFM4) as a new target for the prevention of Alzheimer’s disease.

## Introduction

The apolipoprotein E (*APOE)* gene is among the most studied genes because of the associations of the e4 allele with Alzheimer’s Disease (AD), cognition, and other aging-related traits.^1^ The e4 allele is defined by a specific combination of genotypes of the single nucleotide polymorphisms (SNPs) rs7412 and rs429358. A different combination of genotypes of the same SNPs determines the e2 allele of *APOE* and, interestingly, this allele is associated with extreme human longevity^2^ and delayed onset of cognitive decline.^3^ However, the molecular mechanisms through which the e2 allele influences these traits remains unclear.^4^ To address this knowledge gap, we previously identified a serum protein signature of e2 by correlating *APOE* genotypes of 224 participants of the New England Centenarian Study (NECS)^5^ with 4,137 serum proteins that we profiled using the aptamer-based Somalogic technology.^6^ The analysis identified a signature of 16 serum proteins that highlighted a possible role of e2 in the regulation of inflammation and in lipid regulation via two apolipoproteins in serum: apolipoprotein B (APOB) and apolipoprotein E (APOE). We replicated nine of these associations in 733 plasma samples from the Longenity Gene Project (LGP)^7^ that used the same Somalogic technology.^8^ However, a well-known limitation of aptamer-based technology is the lack of specificty of some of the aptamer binding^9^ and replication using the same technology is insufficient to validate the identity of proteins in the signature. In this paper, we therefore used additional serum proteomics technologies and blood transcriptomics to validate and expand the original signature.

## Results

### Mass spectrometry identifies correlation of *APOE* genotypes with several apolipoproteins

We developed a nano liquid chromatography tandem mass spectrometry (nLC-MS/MS) workflow for in-depth serum proteomic analysis.^10^ We applied this workflow to analyze serum samples from 50 NECS participants representing the *APOE* genotypes e2e2, e2e3, e3e3 and e3e4. Participants’ ages at the time of the blood draw ranged between 54 and 109 years and other characteristics are summarized in **Table 1**. We profiled the samples in triplicate and in batches of 10 samples at a time, with batches including an even representation of *APOE* genotypes and age to reduce confounding of these factors with batch effects. We processed the data with MaxQuant^11^, used ComBat^12^ to remove batch-to-batch variations and, after QC, we identified 398 proteins from 2,654 peptides (**Supplement Table 1**).

**Table 1.** Characteristics on NECS participants included in the study.

|  | *APOE* genotypes |  |  |  |  |
| --- | --- | --- | --- | --- | --- |
|  | e2e2 | e2e3 | e3e3 | e3e4 | Overall |
| N | 7 | 13 | 20 | 10 | 50 |
| Female Gender, N (%) | 5 (71.4%) | 7 (53.8%) | 11 (55.0%) | 5 (50.0%) | 28 (56.0%) |
| Age at blood draw in years<br>Mean (SD) | 70.4 (11.2) | 82.0 (17.5) | 73.5 (12.7) | 80.3 (16.3) | 76.6 (14.9) |
| Years of education Mean (SD) | 14.7 (2.29) | 13.8 (3.66) | 15.4 (3.45) | 16.5 (3.95) | 15.1 (3.51) |

Of the 398 proteins identified in our analysis, 266 overlapped with those included in the Somalogic array used in the original study.^6^ We analyzed the log2-transformed, nLC-MS/MS-based abundance data using linear regression with generalized estimating equations and identified 18 proteins as significantly associated with APOE2 at a 1% false discovery rate (FDR) (**Table 2**) and 39 proteins at 5% FDR (**Supplement Table 1**). The list of 18 proteins included APOB that validated the original result^6^, and APOE that exhibited significant associations with the *APOE* genotypes but with different direction of effects (**Fig 1**).

**Fig 1:**
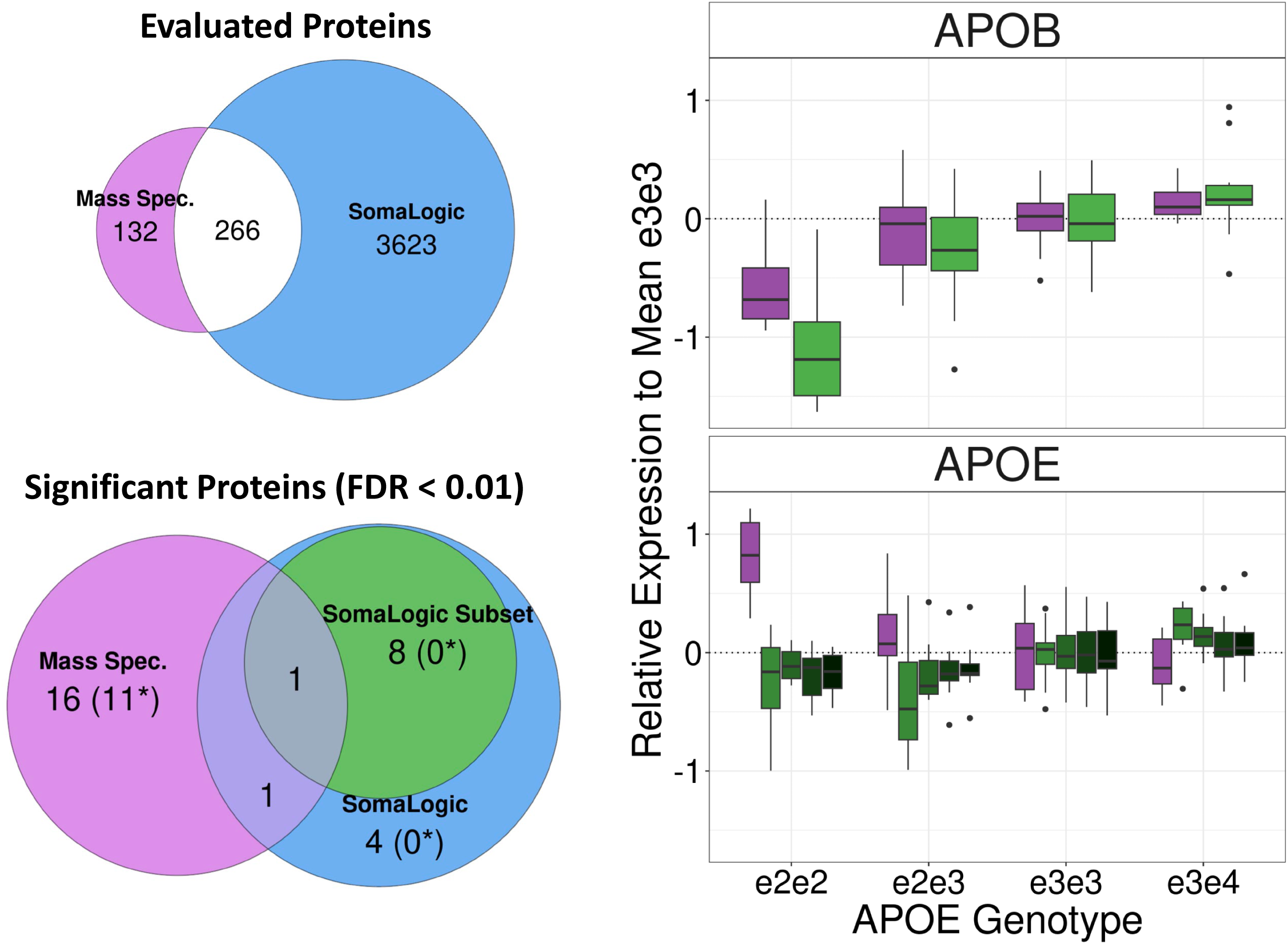
**Top left**: We detected 398 proteins in 50 serum samples with nLC-MS/MS, compared to 3,889 proteins detected with the Somalogic platform and 266 proteins were in common. **Bottom Left**: the Venn diagram displays the number of proteins associated with *APOE* genotypes in the nLC-MS/MS analysis (purple), and in Somalogic-based analysis of the same 50 samples (green). **Right**: Both analyses detected APOB and APOE associations with *APOE* genotypes (purple: nLC-MS/MS data; green Somalogic).

**Table 2.** Association of serum proteins with APOE genotypes in the nLC-MS/MS-based analysis. Columns e2e2, e2e3, and e3e4 represent fold change of expression in carriers of the column genotype relative to e3e3 carriers. Adjusted p: Global P-values adjusted for multiple testing using Benjamini-Hochberg false discovery rate.

| Uniprot | Gene | N peptides | e2e2 | e2e3 | e3e4 | Adjusted P |
| --- | --- | --- | --- | --- | --- | --- |
| P02649 | APOE | 18 | 1.70 | 1.12 | 0.94 | 5.00E-07 |
| P49913 | CAMP | 2 | 1.87 | 1.12 | 0.90 | 1.83E-06 |
| P04114 | APOB | 188 | 0.66 | 0.97 | 1.13 | 1.31E-05 |
| O14791 | APOL1 | 7 | 1.40 | 1.20 | 1.04 | 0.000206 |
| P02654 | APOC1 | 3 | 1.33 | 1.26 | 0.95 | 0.000535 |
| P61626 | LYZ | 3 | 1.30 | 1.15 | 0.85 | 0.000841 |
| P11597 | CETP | 2 | 1.39 | 1.11 | 0.94 | 0.00091 |
| Q01459 | CTBS | 3 | 0.98 | 0.87 | 1.01 | 0.003681 |
| P08185 | SERPINA6 | 10 | 0.92 | 0.81 | 0.92 | 0.004598 |
| P05164 | MPO | 7 | 1.53 | 1.31 | 0.76 | 0.004598 |
| P55056 | APOC4 | 2 | 2.32 | 1.80 | 1.20 | 0.004815 |
| P08311 | CTSG | 3 | 1.62 | 1.41 | 0.73 | 0.004815 |
| Q12841 | FSTL1 | 1 | 1.09 | 0.84 | 0.88 | 0.00624 |
| P59666 | DEFA3 | 2 | 1.63 | 1.32 | 0.81 | 0.007212 |
| P51884 | LUM | 10 | 0.92 | 0.84 | 0.94 | 0.007212 |
| P02774 | GC | 26 | 1.02 | 0.91 | 1.01 | 0.008093 |
| P00747 | PLG | 31 | 1.10 | 0.94 | 0.87 | 0.008788 |
| P01042 | KNG1 | 19 | 1.15 | 0.97 | 1.01 | 0.009867 |

The Somalogic array used in the original study included 23 aptamers linked to 13 apolipoproteins (APOA1, APOA2, APOA5, APOB, APOBC3G, APOC2, APOC3, APOD, APOE, APOF, APOH, APOL1, APOM, APOO). With the exception of APOB and APOE, we did not detect significant associations between these apolipoproteins and the e2 allele in the original analysis.^6^ The nLC-MS/MS-based analysis detected the 12 apolipoproteins APOA4, APOB, APOC1, APOC2, APOC3, APOC4, APOD, APOE, APOF, APOH, APOL1, and APOM in the 50 serum samples. Among these, we detected significant associations between *APOE* genotypes and APOB, APOC1, APOC4, APOE, and APOL1 at 1% FDR, and between *APOE* genotypes and APOC2, APOC3, and APOF at 5% FDR (**Fig 2**). APOC1, APOC2, APOC3, APOC4, APOE, and APOL1 showed increased levels in carriers of the e2 allele: the median fold change of e2e2 carriers compared to e3e3 carriers (FC_22._33) was 1.72, range 1.33 to 2.32, while the median fold change of e2e3 carriers compared to e3e3 carriers (FC_23.33_) was 1.32, range 1.12 to 1.39. Similarly to APOB, the level of APOF decreased with increasing copies of the e2 allele (FC_22.33_= 0.78, FC_23.33_= 0.87, FC_34.33_=0.96, Adj_p=0.03). In addition, we detected a significantly higher level of Cholesteryl ester transfer protein (CETP) in e2e2 carriers compared to e3e3 carriers (FC_22.33_= 1.39, FC_23.33_= 1.11, FC_34.33_=0.94, Adj_p =9.1E-4). The pairwise correlations of these apolipoproteins (**Fig 2**) were stronger in carriers of at least one copy of the e2 allele compared to e3e3 and e3e4 carriers (median correlation for e2e3 carriers = 0.34 vs 0.08 for e3e3 carriers and 0.06 for e3e4 carriers).

**Fig 2:**
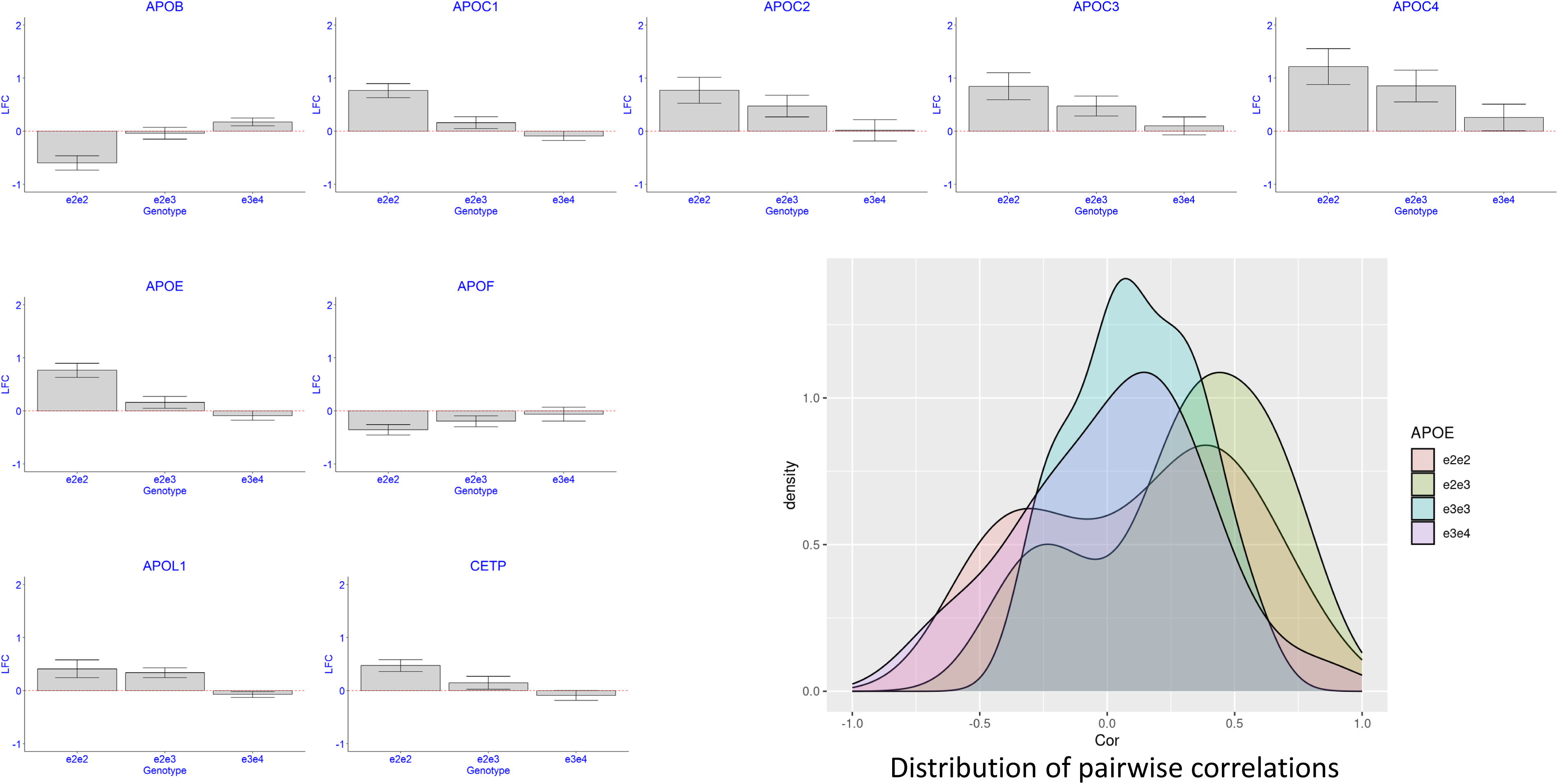
Distribution of Apolipoproteins and CETP by *APOE* genotypes. Bar plots of the estimated log-fold changes of serum proteins comparing e2e2, e2e3 and e3e4 to e3e3 carriers. Bars denote SE. The density plot displays the pairwise correlations between these serum proteins in the four *APOE* genotype groups. Carriers of e2e2 and e2e3 genotype have stronger pairwise correlations.

### Mass spectrometry identifies effects of the e2 allele on several markers of immune response

The nLC-MS/MS-based analysis identified an additional 13 proteins that correlated significantly with the *APOE* genotypes e2e2 and e2e3. Some examples are in **Fig 3**, and the complete list is in **Table 2** **and Supplement Table 1**. This set included Plasminogen (PLG, FC_22.33_= 1.10, FC_23.33_= 0.94, FC_34.33_=0.87, p= 0.008788), and the granule proteins secreted by neutrophils: Cathelicidin antimicrobial peptide (CAMP, FC_22.33_= 1.87, FC_23.33_= 1.12, FC_34.33_=0.90, p=1.8E-6), Cathepsin G (CTSG, FC_22.33_= 1.62, FC_23.33_= 1.41, FC_34.33_=0.73, p= 0.00481456), Defensive Alpha 3 (DEFA3, FC_22.33_= 1.63, FC_23.33_= 1.32, FC_34.33_=0.81, p= 0.0072), and Myeloperoxidase (MPO, FC_22.33_= 1.53, FC_23.33_= 1.31, FC_34.33_=0.76, p= 0.004599). All these proteins were strongly correlated (**Fig 3**), and the magnitude of the pairwise correlations did not change with *APOE* genotypes (median correlation for e2e3 carriers 0.42 vs 0.38 for e3e3 carriers and 0.42 for e3e4 carriers).

**Fig 3:**
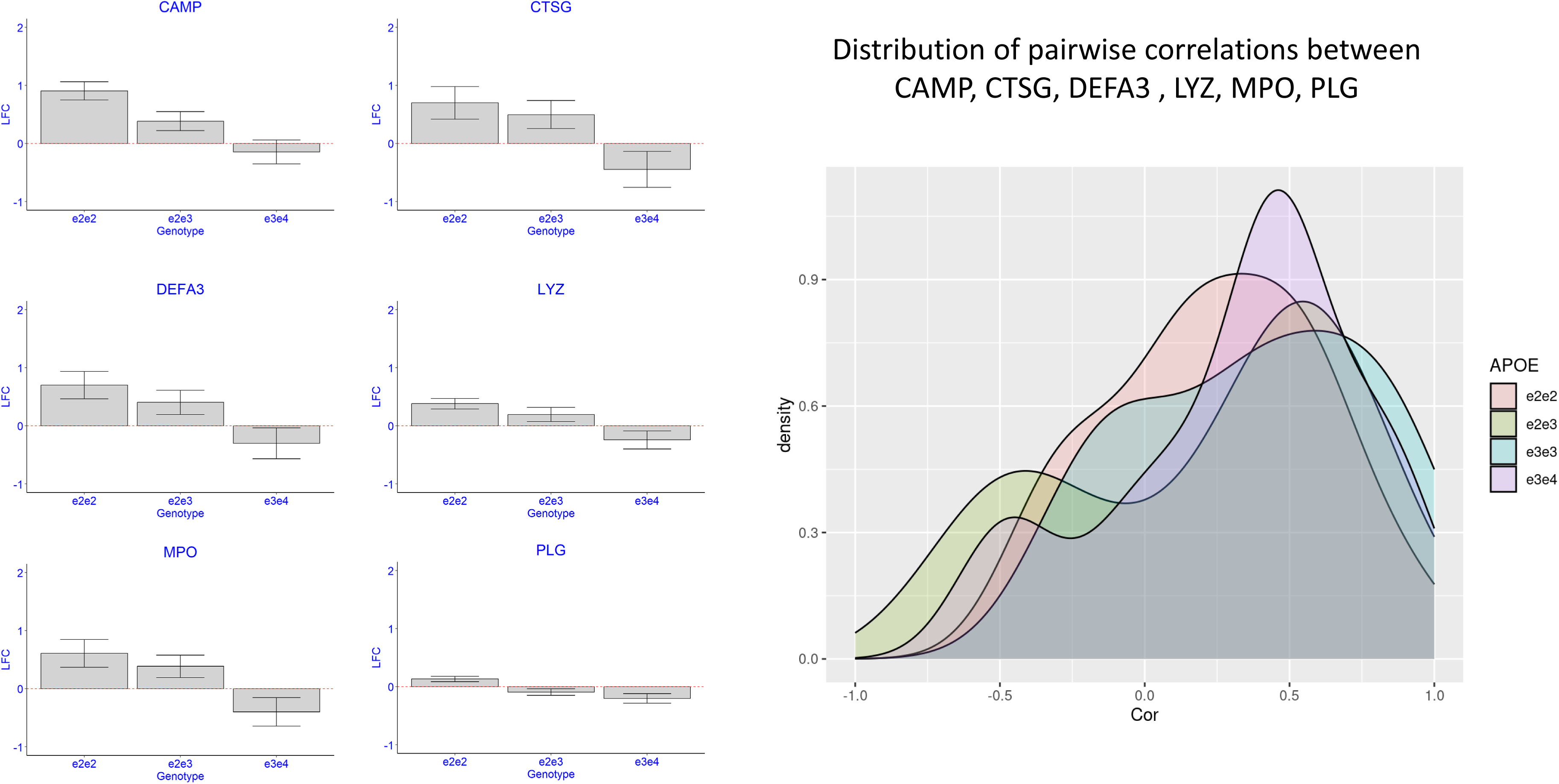
Distribution of inflammatory markers by *APOE* genotypes. Boxplot display the distribution of log-fold change of six inflammatory markers comparing e2e2, e2e3 and e3e4 to e3e3 carriers. The density plot displays the pairwise correlations between these serum proteins in the four *APOE* genotype groups.

### Olink technology validates some of the mass spectrometry-based analysis

We attempted replication of the *APOE* e2-associated proteins in the UK Biobank serum proteomic data, which used the antibody-based Olink Explore 3072 Proximity Extension Assay (PEA).^13^ After QC, we analyzed data from 34,875 participants and for 20 of the 39 proteins that were in the Olink platform. We identified significant associations with consistent effects between the *APOE* genotypes e2e2 and e2e3 and APOC1 (FC_22.33_ =1.21, p=2.34E-29; FC_23.33_ =1.05, p=1.13E-28), APOE (FC_22.33_ =2.23, p=1.3E-177; FC_23.33_ =1.20, p=1.30E-118), FSTL1 (FC_22.33_ =1.03, p=9.99E-3), and lipoprotein(a) (LPA, FC_22.33_ =0.60, p=8.62E-10). However, associations between *APOE* e2e2 genotypes and coagulation factor 11 (F11, p=1.48E-2), vitamin D binding protein (GC, p=2.64E-2) were inconsistent (**Supplement Figure 1**).

### Replication in blood transcriptomics suggests that the e2 allele affects immune response

Except for APOB and APOE, none of the other proteins in the original Somascan-based signature (Supplement Table 3) could be detected with the nLC-MS/MS workflow, thus leaving the issue of cross-platform validation unresolved. We hypothesized that since some of the proteins associated with e2 are markers of inflammatory response, we should be able to observe some of these patterns in blood transcriptomics. To test this hypothesis, we analyzed RNA-seq-based whole-blood transcriptional profiles from 1,348 participants in the Long Life Family Study (LLFS). The ages of the participants ranged from 24 to 107 years, and the data set included seven e2e2 and 203 carriers of e2e3. We correlated ~11K transcripts with *APOE* genotypes using standard linear regression adjusted by sex, and medication for hypertension, type 2 diabetes, high cholesterol, and heart disease. This analysis identified 74 transcripts associated with the e2 allele (either e2e2 or e2e3 genotype) at 5% FDR, and only one transcript associated with the e2e2 genotype at 20% FDR (**Supplement Table 2**). The estimates of the *APOE* genotype effects on gene expression in LLFS blood transcriptomics were positively correlated with the estimates from nLC-MS/MS-based serum proteomics, and we observed good concordance for genes APOL1, LYZ, MPO, and CAMP (**Fig 4a**). The correlation between *APOE* genotype effect estimates on LLFS blood transcriptomics and Somalogic-based serum proteomics was weaker by comparison, and only significant in e2e2 carriers. However, we did observe concordance of the *APOE*-associated patterns of baculoviral IAP repeat containing 2 (BIRC2), leucine rich repeat neuronal 1 (LRRN1), ubiquitin like modifier activating enzyme 2 (UBA2), proteasome activator subunit 1 (PSME1), and VPS29 retromer complex component (VPS29) in e2e3 carriers (**Fig 4b**). In this analysis, we did not adjust by age because of the correlation between the e2 allele and longevity. We also conducted a sensitivity analysis in which we adjusted the results by age and the results were very robust (Supplement Table 3).

**Fig 4:**
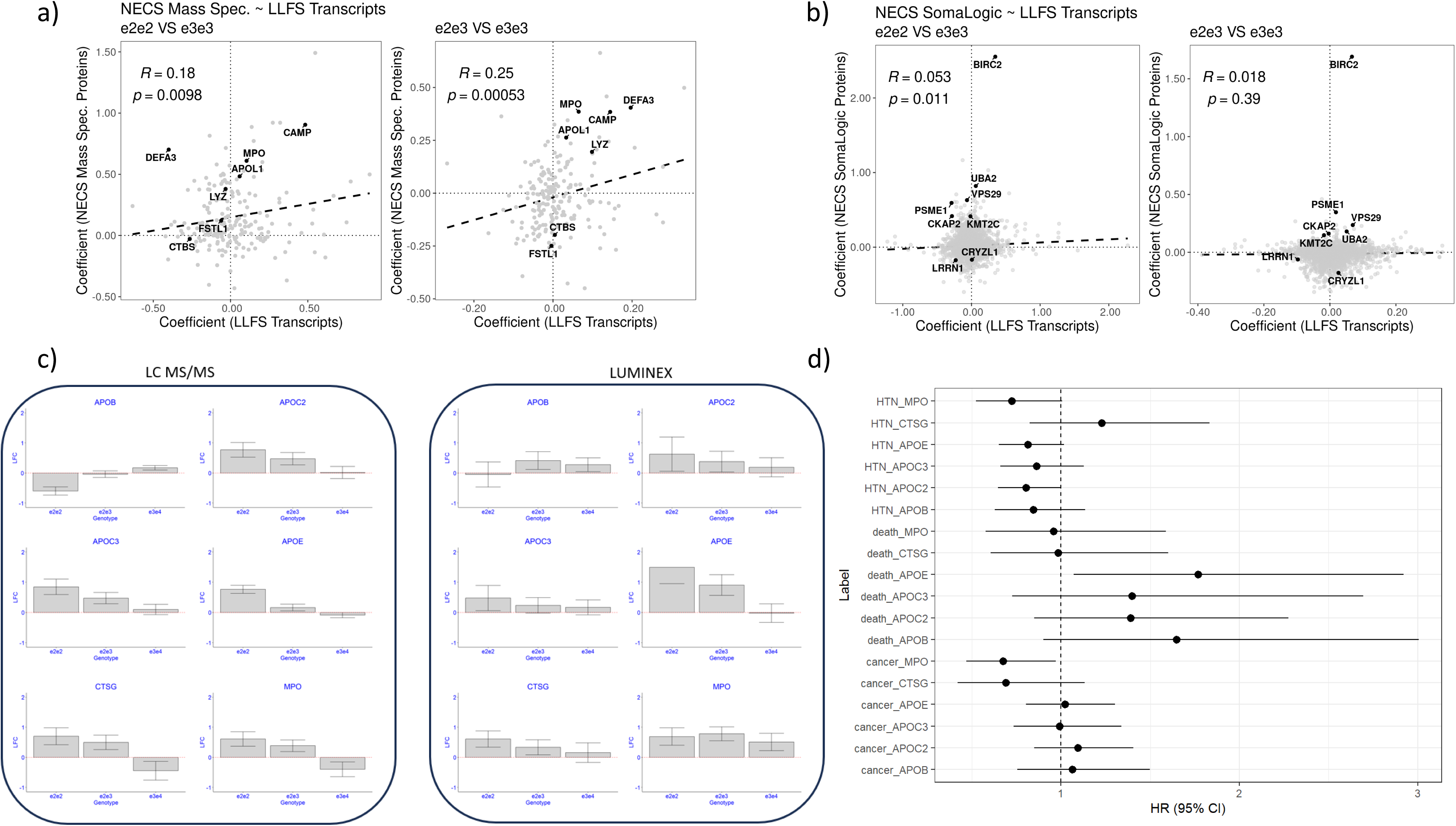
**a)** Correlation of *APOE* effects with gene expression in 1348 blood samples from the LLFS (x-axis), and serum proteins measured with nLC-MS in 50 NECS participants (y-axis). Each dot represents one of the 196 genes that we could map succesfully between the two datasets. **Left**: effects of e2e2 relative to e3e3. **Right**: effects of e2e3 relative to e3e3. **b)** Correlation of *APOE* effects with gene expression in 1348 blood samples from the LLFS (x-axis), and serum proteins measured with Somascan in 220 NECS participants (y-axis). Each dot represents one of the 2023 genes that we could map succesfully between the two datasets. **Left**: effects of e2e2 relative to e3e3. **Right**: effects of e2e3 relative to e3e3. **c)** replication of some of the effects detected with the LC MS/MS analysis and Luminex. **d)** Estimated hazard ratio and 95% confidence intervals for hypertension (HTN), death and cancer for 1 log-fold change of APOB, APOC2, APOC3, APOE, CTSG and MPO.

### *APOE* genotype targets predict aging traits

We measured serum levels of APOB, APOC2, APOC3, APOE, CTSG, and MPO in 106 NECS participants, including four e2e2 carriers and 26 e2e3 carriers, using a combination of Luminex and ELISA proteomics platforms. This set also included 59 samples repeated over an average of 16 years. We analyzed the repeated measures of these 6 biomarkers to (1) replicate their associations with *APOE* genotypes, (2) identify the biomarkers that change over time, and (3) assess whether these biomarkers predict the onset of aging related diseases and cognitive decline, as measured by the Telephone Interview for Cognitive Status (TICS). The analysis validated the associations of APOE, CTSG, and MPO with *APOE* genotypes (**Fig 4**, **Table 3**). While the associations between serum levels of APOB, APOC2, and APOC3 had concordant effects, they did not reach statistical significance. Overall, the analysis also showed that levels of APOC2, APOC3, and MPO declined significantly over time. Levels of APOB and APOE also showed a negative trend over time, but the associations failed to reach statistical significance Additionally, higher CTSG levels were positively correlated with a faster rate of cognitive decline as measured by TICS (beta= 0.08, p=0.02) and we observed a similar but non-significant trend for MPO. Time to event analysis showed that higher levels of MPO decreased the risk of all cancers combined (Hazard ratio for 1-fold change of serum level: HR=0.67, p=0.033) and hypertension (HR=0.72, p=0.05) (**Fig 4c, Table 4**). Furthermore, higher levels of APOC2 were associated with decreased hazard for hypertension (HR=0.80, p=0.047). In contrast, higher levels of APOE predicted a significant 77% increase in risk for death (HR= 1.77, p=0.026). Similarly, higher levels of APOB, APOC2, and APOC3 were positively correlated with increased risk of death although those associations were not statistically significant.

**Table 3.**
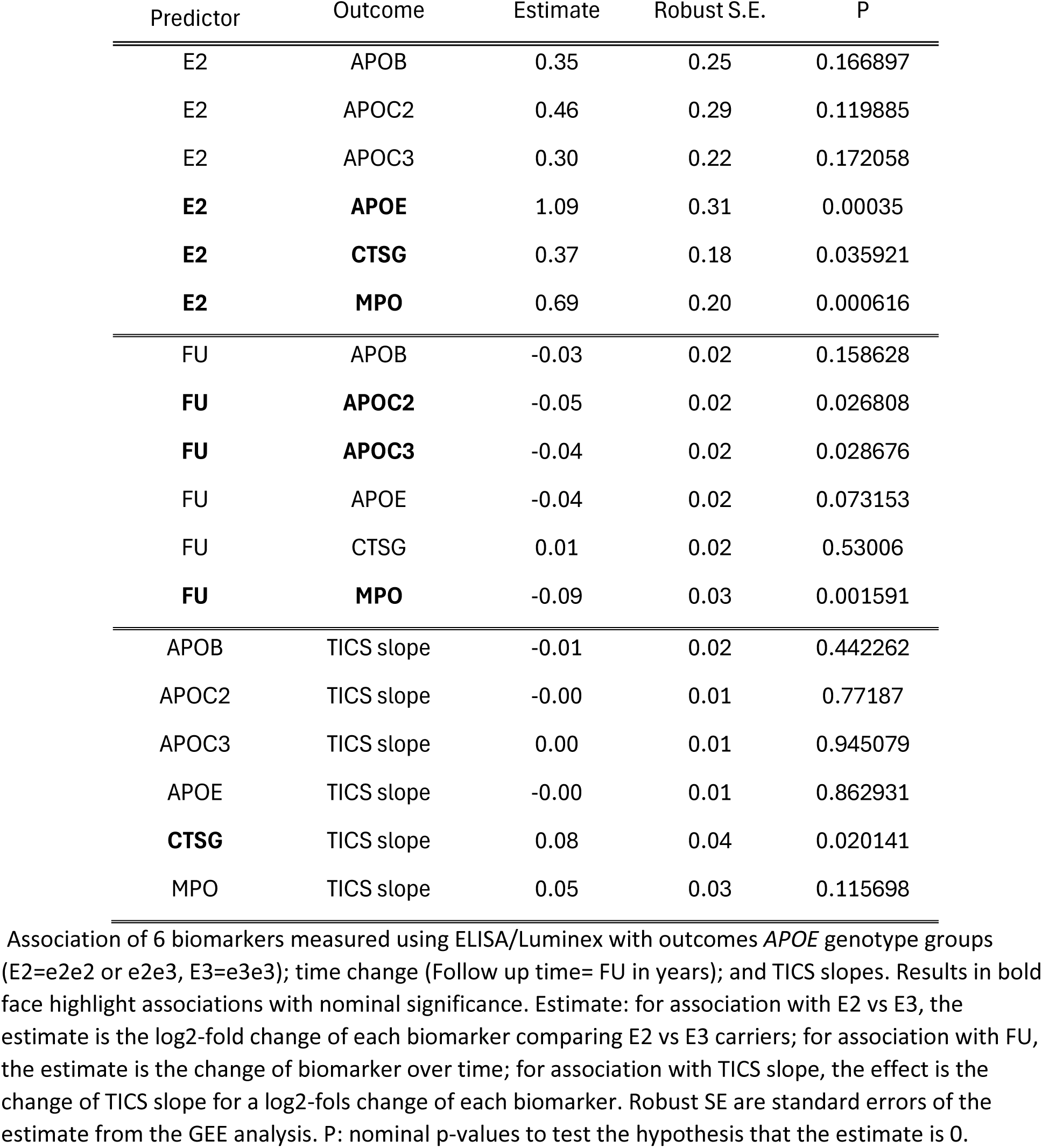
Association of 6 biomarkers measured using ELISA/Luminex with outcomes *APOE* genotype groups (E2=e2e2 or e2e3, E3=e3e3); time change (Follow up time= FU in years); and TICS slopes. Results in bold face highlight associations with nominal significance. Estimate: for association with E2 vs E3, the estimate is the log2-fold change of each biomarker comparing E2 vs E3 carriers; for association with FU, the estimate is the change of biomarker over time; for association with TICS slope, the effect is the change of TICS slope for a log2-fols change of each biomarker. Robust SE are standard errors of the estimate from the GEE analysis. P: nominal p-values to test the hypothesis that the estimate is 0.

## Discussion

We validated and expanded a serum protein signature of *APOE* genotypes using a combination of mass-spectrometry, ELISA, Luminex, and antibody-based Olink PAE proteomics, as well as blood transcriptomics. We replicated the association between APOB and the e2 allele of *APOE*. We corrected the pattern of association between *APOE* genotypes and serum level of APOE, and we detected new associations between *APOE* genotypes and the complex of apolipoproteins APOC1, APOC4, APOC2, APOC3, APOE, APOF and APOL1. In addition, we discovered 13 new proteins that significantly correlated with *APOE* genotypes and included a signature of granule proteins secreted from neutrophils (CAMP, CTSG, DEFA3, and MPO).

### Role of APOE2 in lipids regulation

The role of APOE in lipids regulation is well studied^14^, and both our work and that of others have shown that the e2 allele correlates with the distribution of a variety of lipid species.^15–17^ This new analysis confirms that carriers of the e2e2 and e2e3 genotypes have lower levels of APOB and APOF compared to carriers of e3e3. The increased level of APOB in the serum of e3 or e4 carriers is well established^6, 18^, and a high level of APOB is a validated biomarker of cardiovascular disease risk.^19^ Lower levels of APOB in carriers of e2e2 and e2e3 genotypes compared to carriers of e3e3 are also established^20^, although the mechanism of action remains unclear.^21^ The decreased levels of APOB we observed in e2 carriers may result from reduced production of very low density lipoprotein VLDL and increased clearance of remnant particles. The low affinity of the e2 allele of *APOE* for low density lipoprotein receptor (LDLR) may redirect lipoproteins to alternative clearance pathways involving low-density lipoprotein receptor-related protein 1 (LRP1) and heparan sulfate proteoglycan (HSPG). This hypothesis is consistent with the increased levels of APOC1-4 and APOE we observed in our data. The simultaneously low level of APOF and high level of CETP in e2 carriers that we observed is consistent with evidence that APOF is a natural inhibitor of CETP and a key regulator of lipoprotein metabolism.^22^ Our data suggests that the strengths of coregulation between APOF and CETP changes with *APOE* genotypes: we observed a 23% negative correlation in e2e2 carriers, a 12% negative correlation in e2e3 carriers, a 41% negative correlation in e3e4 carriers and a positive correlation in e3e3 carriers (**Supplement Figure 1**).

Our analysis detected a significant increase in the serum level of APOE, APOC1, APOC2, APOC3 and APOC4 in carriers of one e2 allele compared to e3e3 carriers. In addition, many apolipoproteins showed stronger correlations in e2 carriers compared to e3e3 carriers. APOE, APOC1, APOC2, and APOC4 are part of a gene cluster on chromosome 19 that are under control of the same cis-regulatory elements, and their coordinated changes likely reflect this co-regulation. APOC3 is on chromosome 11, and we observed a stronger correlation of this apolipoprotein with the complex APOE, APOC1, APOC2 and APOC4 in e2e2 and e2e3 carriers (correlation ranging between 0.55 and 0.82 in both e2e2 and e2e3 carriers) than e4 carriers (**Fig 2**). Lower cerebrospinal fluid levels of APOC1 in carriers of e4 compared to e3 have been reported previously.^23^ This protein plays a role in the metabolism of lipoproteins, but the role is unclear.^24^ APOC2 and APOC3 regulate plasma cholesterol and triglycerides. APOC2 specifically activates lipoprotein lipase that breaks down triglycerides to provide free fatty acids for cells. While some of these changes in e2 carriers are consistent with a better health profiles, increasing levels of APOC3 have been associated with increasing risk for cardiovascular disease, and inhibition of APOC3 has been suggested as a possible therapeutic target.^25^ Additional analyses are necessary, but considering the elevated level of APOC3 in e2 carriers, our results suggest that the elevated APOC3 level in patients with metabolic disorder could be a compensatory mechanism rather than a causative mechanism of the disease. The results are also consistent with a multi-omics analysis showing that the *APOE* genotype significantly impacts bioenergetic pathways and metabolic associations, particularly in relation to inflammatory markers and insulin sensitivity.^26^

The association between *APOE* genotypes and serum levels of APOE in the previous Somascan-based analysis and the new analyses was discordant. This discrepancy may be due to a lack of specificity for the aptamers that target APOE. Our new analyses correct the previous finding and are more consistent with recent findings showing that higher levels of APOE in plasma were associated with better cognitive performance.^27^ The association between *APOE* genotypes and serum level of APOE was also consistent with the recent results from the UK biobank proteome study.^13^

### *APOE* alleles affect inflammatory markers

The agreement between the gene signature in blood and serum suggests that some of the e2-associated mechanisms may be mediated by inflammation. However, it is not clear if the role of e2 in regulating inflammatory processes is protective. Recent studies have indeed raised concerns about the pleiotropic effect of e2e2 that warrant further investigations.^28–30^ CAMP is an antimicrobial protein linked to the innate immune system with documented role in atherosclerosis.^31^ In our analysis, e2e2 carriers have 87% increased level of CAMP compared to e3e3 carriers (p_adj=1.83E-6). MPO (Table 2) is a heme protein synthesized during myeloid differentiation and an integral component of the innate immune system but it is also responsible for increased tissue damage during inflammation.^32^ MPO has emerged as a potential target for cardiovascular disease, and the accumulation of MPO in brain tissue has been linked to AD.^33^ PLG dissolves the fibrins of blood clots, but can also influence immune response^34^. These results support the role of e2 and e2e2 in modulating inflammatory response, consistent with the inclusion of BIRC2 and LRRN1 in our original protein signature^6^. In addition to CAMP and MPO, e2e2 carriers had elevated levels of lysozyme (LYZ), CTSG, DEFA3, CAMP, MPO, LYZ, CTSG, DEFA3. Neutrophils release all these proteins, and their high levels in serum may modulate the activation of innate immune cells.^35^ The correlation between *APOE* allele and this signature suggests that the role of e2 in modulating inflammatory response could be enacted in non-e2 carriers by direct targeting of this signature.

Interestingly, the work by Vandenberghe-Dürr S et al^36^ showed that inhibition of olfactomedin 4 (OLFM4) also promotes bacterial clearance by neutrophils.^36^ As a result, we conjecture that OLFM4 may be an interesting target for AD treatment or healthy cognitive aging. OLFM4 is a well-studied marker of many cancers, and the inhibition of this target should be well characterized.

### Robust validation of proteomics findings requires multiple approaches

Serum proteomics is a very popular approach for biomarker discovery and for understanding disease mechanisms. However, our results underscore that no single proteomics platform is sufficient to draw robust biological conclusions. Validation through multiple technologies and integration of diverse omics layers is essential for accurate interpretation and reproducibility of findings.

## Supplement Methods

### Study population

#### New England Centenarians (NECS)

We used serum samples from blood from NECS participants^5^ who included centenarians, centenarians’ offspring, and subjects without familial longevity. Participants are followed longitudinally to update their medical history and their cognitive status using the Telephone Interview for Cognitive State (TICS)^37^. All samples were frozen on arrival to the molecular lab and maintained at −80°C until assays were performed. The 50 samples used for serum proteomics with LC-MS were a subset of the 224 samples originally profiled with the Somascan technology^10^. Their ages ranged from 50 to 100 years. The set included 7 e2e2 carriers (Table 1).

#### Long Life Family Study (LLFS)

Whole blood RNA sequencing transcriptomic and genotype data were obtained from the LLFS, a family-based study of healthy aging and longevity^38^. These subjects’ ages ranged from 24 to 107, with a mean age of 69.1 years at the time of the blood draw. Transcriptomic profiling was performed in 30 separate batches, with the number of subjects profiled per batch ranging from 23 to 82.

### Serum proteomics

#### nLC-MS/MS Workflow

We used a labelled LC-MS/MS workflow with 11 tandem mass tags (TMT). We profiled samples in five pools of ten samples each and an 11^th^ channel containing a reference standard as a mixture of all 50 serum samples. Each pool included similar distributions of *APOE* genotypes. Each pool was run in triplicate, resulting in a total of 150 sample profiles and 15 control samples. Full details of sample preparation are described in reference^39^.

#### Processing of Mass Spectrometry Data

We used MaxQuant 1.6.17 for peptide quantification.^40^ Filtration criteria for protein matches included 1% false discovery rate, and ≥ 1 unique peptide resulting in a filtered set of 11,584 peptides across 1,473 proteins. We removed 461 peptides associated with 12 depleted proteins (ALBU, APOA1, APOA2, CRP, A1AG1, A1AG2, A1AT, A2MG, FIB, HPT, IGH, TRFE) and a single peptide associated with UniProt identifiers S4R460, which had been removed from the UniProt database. This filtering step produced 11,122 peptides mapping to 1,450 proteins. Next, we filtered peptides based on missingness. First, we removed peptides with missingness across all profiles, resulting in the removal of 743 peptides. Next, we removed additional peptides with missingness in at least 20% of profiles, or missingness in at least 20% of batches, or missingness in at least 20% of pools, i.e. 1 out of 5. Of the 11,122 assigned peptides, 8,469 were removed based on all missingness criteria, resulting in 2,653 peptides in 398 proteins for subsequent analyses. We aggregated measurements of protein expression by summing the associated peptides. Prior to the aggregation, missing peptide values were imputed by drawing from a uniform distribution with a range of 0 to the minimum peptide measurement of each batch. Each profile was then normalized by dividing expression levels by their respective 10% trimmed mean, followed by a log2-transformation. Finally, the normalized profiles were batch corrected to reduce the impact of technical variability using ComBat (v3.44.0).^41^

#### Processing Of Luminex Data

Biomarkers were detected and quantified using both enzyme-linked immunosorbent assays (ELISA) [Cathepsin G], 3-Plex [MPO]; and EMD Millipore Human Apolipoprotein [ApoB, ApoE, Apo C2, and Apo C3] according to each manufacturers’ protocol. All Luminex assays were miniaturized using 96-well DropArray plates and the LT210 plate washer (Curiox Biosystems, Inc), with quantitation performed on a Luminex Magpix instrument equipped with xPONENT 4.2 software to fit standard curves. The commercial ELISA assays used (Bio-Techne Human Cathepsin G) were measured on a SpectraMax i3x Plate Reader (Molecular Devices, LLC) at a wavelength of 450 nm. SoftMax Pro 7.0.3 software was used to fit all standard curves.

### Statistical analysis

#### Statistical Analysis of Mass Spectrometry Data

We evaluated the differences in log2-protein expression between the four *APOE* genotypes, e3e3, e2e2, e2e3, and e3e4 using linear regression adjusting for age at blood draw, year-of-collection, and gender. We used generalized estimating equations (GEE) to estimate the regression coefficients and robust standard errors and account for within-sample variability of each triplicate (geepack, v1.3.4). We assessed the global differences between genotypes using the log-likelihood ratio chi-square tests with 3 degrees of freedom. We corrected p-values for multiple hypothesis testing using the Benjamini-Hochberg False Discovery Rate (FDR) correction.^42^

#### Statistical Analysis of SomaScan Data

The SomaScan data were included in reference^6^ and comprised 4,785 aptamers mapping to 4,118 proteins. We removed 147 aptamers no longer included in more recent versions and updated UniProt IDs, further removing 233 aptamers mapping to mouse protein, Q99LC4, and updating an additional 36 aptamers. The filtered data comprised 4,405 aptamers in 3,889 proteins, and only 266 proteins (353 aptamers) were shared with the processed mass spectrometry data, while 3,623 proteins (4,052 aptamers) were detected only in the SomaScan data, and 132 proteins were detected only in the mass spectrometry data. We re-analyzed the subset of SomaScan data comprising the same 50 subjects profiled with mass spectrometry following the same procedure as the published SomaScan study.^6^ Briefly, protein RFU were log2-transformed and values beyond three standard deviations of the 5% trimmed mean were removed. We next analyzed the differences in the log2-protein expression between the four *APOE* genotypes e3e3, e2e2, e2e3, and e3e4, using linear regression. We tested for global differences between *APOE* genotypes using ANOVA. We corrected p-values for multiple hypothesis testing using the Benjamini-Hochberg False Discovery Rate (FDR) correction.^42^ We also updated the multiple testing correction of the results from the original publication^6^ following the update of their aptamer annotation.

#### Statistical analysis of Luminex and ELISA Measurements

We correlated the repeated measurements of the six proteins with *APOE* genotype groups (E2=e2e2 or e2e3, E3=e3e3, E4=e3e4), age at blood draw, follow up time, sex, and education using linear regression of log2-transformed raw values. We estimated regression coefficients and robust standard errors using GEE with the geepack, v1.3.4. We used mixed effect models with random intercept and random slope in 109 NECS participants to estimate individual rate of change of the TICS score assessed at least 2 times. The estimated slopes were used as outcomes of linear regression to assess the effect of these six biomarkers of the rate of change of cognitive function. We estimated the effect of these six biomarkers on mortality and incident events of cancer, cardiovascular disease, type 2 diabetes, hypertension, and stroke using Cox proportional hazard regression, adjusted for age at blood draw, sex, and years of education. The analyses were stratified by generation, defined as birth year earlier than 1935 to address non-proportional hazards of the generations of centenarians and their offspring. Repeated measurements of the six biomarkers were averaged before these analyses.

#### Statistical Analysis of Transcriptomic Data

The RNAseq data was generated from March 2021 to November 2023 at Washington University in St. Louis. Details are provided in reference.^43^ Briefly, total RNA was extracted from PAXgene™ Blood RNA tubes using the Qiagen PreAnalytiX PAXgene Blood miRNA Kit (Qiagen, Valencia, CA). The Qiagen QIAcube extraction robot performed the extraction according to the company’s protocol. The RNA-Seq data were processed with the nf-core/RNASeq pipeline version 3.3 using STAR/RSEM (https://zenodo.org/records/5146005). Samples with high intergenic read percentage (> 8%), suspicious sex chromosome gene expression, and without sufficient genotype or phenotype information were excluded. Transcripts with less than 10 counts per million in at least 3% of samples across the original 4189-sample series were excluded, yielding a final set of 11,229 transcripts for the analysis. Raw transcripts counts were then normalized using the R package *DESeq2* and log2-transformed. We analyzed the effects of *APOE* genotypes on gene expression at the baseline visit, using linear regression of the log2-transformed normalized expression data. We minimally adjusted the analysis by medication for hypertension, type 2 diabetes, dyslipidemia, and heart disease, and repeated the analysis adjusting also for age.

#### Statistical analysis of the UK Biobank data

We correlated *APOE* genotypes from participants in the UK Biobank with Olink proteomics data (log2-transformed, normalized expression) at baseline (N=53,013).^13^ We excluded participants and proteins with more than 10% missing values from the dataset (N=34,875) and imputed the remaining missing values using the median of each analyte. Then, we applied a linear regression model to correlate the list of 39 proteins associated with the *APOE* genotypes, adjusting for age, sex, and four genome-wide principal components of the genetic data.

All analyses were conducted using the R software. The analyses of the UK Biobank data were conducted in the Research Analysis Platform (https://ukbiobank.dnanexus.com).

## Data availability

Both serum proteomics data in the NECS and blood transcriptomics in the LLFS are available from the ELITE portal (https://eliteportal.synapse.org/)

## Supplement figures

**Supplement Figure 1:**
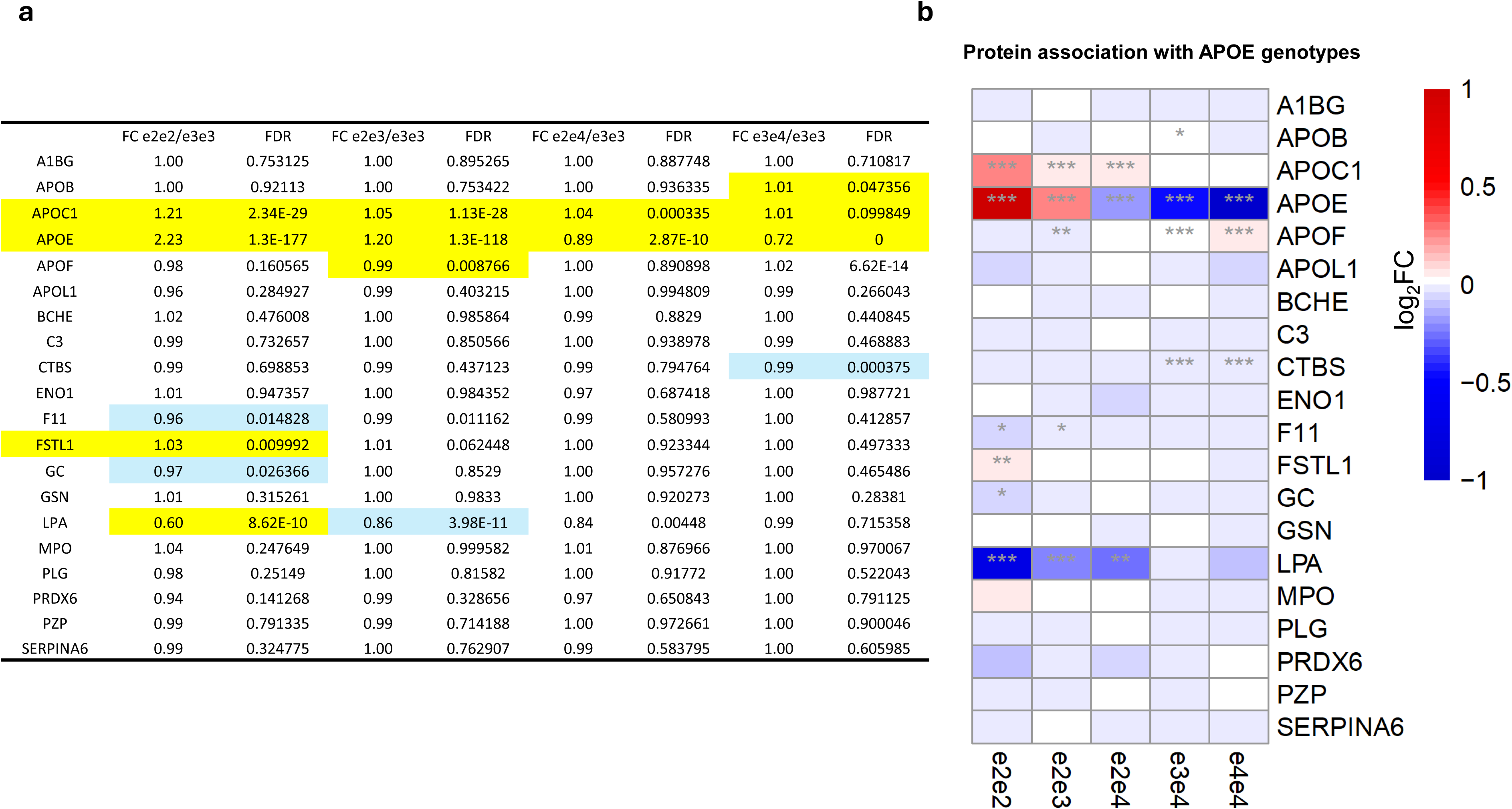
Protein association with APOE genotypes in the UK Biobank. A.FC=fold changes. FDR= false discovery rate. b. Linear regression analysis on all participants. The colors in the heatmap denote the regression coefficients (log_2_FC) for APOE genotypes (blue:log_2_FC<0, white: log_2_FC=0, red: log_2_FC>0), and asterisk symbols imply statistical significance: *FDR<0.05, **FDR<0.01, and ***FDR<0.001.

## References

1. Belloy ME, Andrews SJ, Le Guen Y, Cuccaro M, Farrer LA, Napolioni V, Greicius MD. APOE Genotype and Alzheimer Disease Risk Across Age, Sex, and Population Ancestry. JAMA Neurology. 2023. doi: 10.1001/jamaneurol.2023.3599.

2. Sebastiani P, Gurinovich A, Nygaard M, Sasaki T, Sweigart B, Bae H, Andersen SL, Villa F, Atzmon G, Christensen K, Arai Y, Barzilai N, Puca A, Christiansen L, Hirose N, Perls TT. APOE Alleles and Extreme Human Longevity. J Gerontol A Biol Sci Med Sci. 2019;74(1):44–51. doi: 10.1093/gerona/gly174. PubMed PMID: 30060062; PMCID: PMC6298189.

3. Sweigart B, Andersen SL, Gurinovich A, Cosentino S, Schupf N, Perls TT, Sebastiani P. APOE E2/E2 Is Associated with Slower Rate of Cognitive Decline with Age. J Alzheimers Dis. 2021. Epub 2021/08/10. doi: 10.3233/JAD-201205. PubMed PMID: 34366332.

4. Toribio-Fernández R, Tristão-Pereira C, Carlos Silla-Castro J, Callejas S, Oliva B, Fernandez-Nueda I, Garcia-Lunar I, Perez-Herreras C, María Ordovás J, Martin P, Blanco-Kelly F, Ayuso C, Lara-Pezzi E, Fernandez-Ortiz A, Garcia-Alvarez A, Dopazo A, Sanchez-Cabo F, Ibanez B, Cortes-Canteli M, Fuster V. Apolipoprotein E-ε2 and Resistance to Atherosclerosis in Midlife: The PESA Observational Study. Circulation Research. 2024;134(4):411–24. doi: 10.1161/CIRCRESAHA.123.323921.

5. Sebastiani P, Perls TT. The Genetics of Extreme Longevity: Lessons from the New England Centenarian Study. Frontiers in Genetics. 2012;3. doi: 10.3389/fgene.2012.00277.

6. Sebastiani P, Monti S, Morris M, Gurinovich A, Toshiko T, Andersen SL, Sweigart B, Ferrucci L, Jennings LL, Glass DJ, Perls TT. A serum protein signature of APOE genotypes in centenarians. Aging Cell. 2019;18(6):e13023. Epub 20190805. doi: 10.1111/acel.13023. PubMed PMID: 31385390; PMCID: PMC6826130.

7. Zhang WB, Aleksic S, Gao T, Weiss EF, Demetriou E, Verghese J, Holtzer R, Barzilai N, Milman S. Insulin-like Growth Factor-1 and IGF Binding Proteins Predict All-Cause Mortality and Morbidity in Older Adults. Cells. 2020;9(6). doi: 10.3390/cells9061368. PubMed PMID: 32492897; PMCID: PMC7349399.

8. Gurinovich A, Song Z, Zhang W, Federico A, Monti S, Andersen SL, Jennings LL, Glass DJ, Barzilai N, Millman S, Perls TT, Sebastiani P. Effect of longevity genetic variants on the molecular aging rate. Geroscience. 2021;43(3):1237–51. Epub 20210504. doi: 10.1007/s11357-021-00376-4. PubMed PMID: 33948810; PMCID: PMC8190315.

9. Haslam DE, Li J, Dillon ST, Gu X, Cao Y, Zeleznik OA, Sasamoto N, Zhang X, Eliassen AH, Liang L, Stampfer MJ, Mora S, Chen Z-Z, Terry KL, Gerszten RE, Hu FB, Chan AT, Libermann TA, Bhupathiraju SN. Stability and reproducibility of proteomic profiles in epidemiological studies: comparing the Olink and SOMAscan platforms. Proteomics. 2022;22(13-14):2100170. doi: 10.1002/pmic.202100170.

10. Sebastiani P, Federico A, Morris M, Gurinovich A, Tanaka T, Chandler KB, Andersen SL, Denis G, Costello CE, Ferrucci L, Jennings L, Glass DJ, Monti S, Perls TT. Protein signatures of centenarians and their offspring suggest centenarians age slower than other humans. Aging Cell. 2021;20(2):e13290. Epub 20210129. doi: 10.1111/acel.13290. PubMed PMID: 33512769; PMCID: PMC7884029.

11. Tyanova S, Temu T, Cox J. The MaxQuant computational platform for mass spectrometry-based shotgun proteomics. Nat Protoc. 2016;11(12):2301–19. doi: 10.1038/nprot.2016.136. PubMed PMID: 27809316.

12. Johnson WE, Li C, Rabinovic A. Adjusting batch effects in microarray expression data using empirical Bayes methods. Biostatistics. 2006;8(1):118–27. doi: 10.1093/biostatistics/kxj037.

13. Sun BB, Chiou J, Traylor M, Benner C, Hsu Y-H, Richardson TG, Surendran P, Mahajan A, Robins C, Vasquez-Grinnell SG, Hou L, Kvikstad EM, Burren OS, Davitte J, Ferber KL, Gillies CE, Hedman ÅK, Hu S, Lin T, Mikkilineni R, Pendergrass RK, Pickering C, Prins B, Baird D, Chen C-Y, Ward LD, Deaton AM, Welsh S, Willis CM, Lehner N, Arnold M, Wörheide MA, Suhre K, Kastenmüller G, Sethi A, Cule M, Raj A, Kang HM, Burkitt-Gray L, Melamud E, Black MH, Fauman EB, Howson JMM, Kang HM, McCarthy MI, Nioi P, Petrovski S, Scott RA, Smith EN, Szalma S, Waterworth DM, Mitnaul LJ, Szustakowski JD, Gibson BW, Miller MR, Whelan CD, Alnylam Human G, AstraZeneca Genomics I, Biogen Biobank T, Bristol Myers S, Genentech Human G, GlaxoSmithKline Genomic S, Pfizer Integrative B, Population Analytics of Janssen Data S, Regeneron Genetics C. Plasma proteomic associations with genetics and health in the UK Biobank. Nature. 2023;622(7982):329–38. doi: 10.1038/s41586-023-06592-6.

14. Liu Z, Li W, Geng L, Sun L, Wang Q, Yu Y, Yan P, Liang C, Ren J, Song M, Zhao Q, Lei J, Cai Y, Li J, Yan K, Wu Z, Chu Q, Li J, Wang S, Li C, Han JJ, Hernandez-Benitez R, Shyh-Chang N, Belmonte JCI, Zhang W, Qu J, Liu GH. Cross-species metabolomic analysis identifies uridine as a potent regeneration promoting factor. Cell Discov. 2022;8(1):6. Epub 20220201. doi: 10.1038/s41421-021-00361-3. PubMed PMID: 35102134; PMCID: PMC8803930.

15. Sebastiani P, Song Z, Ellis D, Tian Q, Schwaiger-Haber M, Stancliffe E, Lustgarten MS, Funk CC, Baloni P, Yao CH, Joshi S, Marron MM, Gurinovich A, Li M, Leshchyk A, Xiang Q, Andersen SL, Feitosa MF, Ukraintseva S, Soerensen M, Fiehn O, Ordovas JM, Haigis M, Monti S, Barzilai N, Milman S, Ferrucci L, Rappaport N, Patti GJ, Perls TT. A metabolomic signature of the APOE2 allele. Geroscience. 2023;45(1):415–26. Epub 2022/08/24. doi: 10.1007/s11357-022-00646-9. PubMed PMID: 35997888; PMCID: PMC9886693.

16. Wang T, Huynh K, Giles C, Mellett NA, Duong T, Nguyen A, Lim WLF, Smith AA, Olshansky G, Cadby G, Hung J, Hui J, Beilby J, Watts GF, Chatterjee P, Martins I, Laws SM, Bush AI, Rowe CC, Villemagne VL, Ames D, Masters CL, Taddei K, Dore V, Fripp J, Arnold M, Kastenmuller G, Nho K, Saykin AJ, Baillie R, Han X, Martins RN, Moses EK, Kaddurah-Daouk R, Meikle PJ. APOE epsilon2 resilience for Alzheimer’s disease is mediated by plasma lipid species: Analysis of three independent cohort studies. Alzheimers Dement. 2022. Epub 2022/01/26. doi: 10.1002/alz.12538. PubMed PMID: 35077012.

17. Sebastiani P, Monti S, Lustgarten MS, Song Z, Ellis D, Tian Q, Schwaiger-Haber M, Stancliffe E, Leshchyk A, Short MI, Ardisson Korat AV, Gurinovich A, Karagiannis T, Li M, Lords HJ, Xiang Q, Marron MM, Bae H, Feitosa MF, Wojczynski MK, O’Connell JR, Montasser ME, Schupf N, Arbeev K, Yashin A, Schork N, Christensen K, Andersen SL, Ferrucci L, Rappaport N, Perls TT, Patti GJ. Metabolite signatures of chronological age, aging, survival, and longevity. Cell Rep. 2024;43(11):114913. Epub 20241105. doi: 10.1016/j.celrep.2024.114913. PubMed PMID: 39504246.

18. Welty FK, Lichtenstein AH, Barrett PHR, Jenner JL, Dolnikowski GG, Schaefer EJ. Effects of ApoE Genotype on ApoB-48 and ApoB-100 Kinetics With Stable Isotopes in Humans. Arteriosclerosis, Thrombosis, and Vascular Biology. 2000;20(7):1807–10. doi: 10.1161/01.ATV.20.7.1807.

19. Welty FK, Lahoz C, Tucker KL, Ordovas JM, Wilson PWF, Schaefer EJ. Frequency of ApoB and ApoE Gene Mutations as Causes of Hypobetalipoproteinemia in the Framingham Offspring Population. Arteriosclerosis, Thrombosis, and Vascular Biology. 1998;18(11):1745–51. doi: 10.1161/01.ATV.18.11.1745.

20. Blanchard V, Ramin-Mangata S, Billon-Crossouard S, Aguesse A, Durand M, Chemello K, Nativel B, Flet L, Chétiveaux M, Jacobi D, Bard J-M, Ouguerram K, Lambert G, Krempf M, Croyal M. Kinetics of plasma apolipoprotein E isoforms by LC-MS/MS: a pilot study. Journal of Lipid Research. 2018;59(5):892–900. doi: 10.1194/jlr.P083576.

21. Khalil YA, Rabès J-P, Boileau C, Varret M. APOE gene variants in primary dyslipidemia. Atherosclerosis. 2021;328:11–22. doi: 10.1016/j.atherosclerosis.2021.05.007.

22. Liu Y, Morton RE. Apolipoprotein F: a natural inhibitor of cholesteryl ester transfer protein and a key regulator of lipoprotein metabolism. Curr Opin Lipidol. 2020;31(4):194–9. doi: 10.1097/mol.0000000000000688. PubMed PMID: 32520778; PMCID: PMC8020876.

23. Cudaback E, Li X, Yang Y, Yoo T, Montine KS, Craft S, Montine TJ, Keene CD. Apolipoprotein C-I is an APOE genotype-dependent suppressor of glial activation. J Neuroinflammation. 2012;9:192. Epub 20120810. doi: 10.1186/1742-2094-9-192. PubMed PMID: 22883744; PMCID: PMC3490924.

24. Rouland A, Masson D, Lagrost L, Vergès B, Gautier T, Bouillet B. Role of apolipoprotein C1 in lipoprotein metabolism, atherosclerosis and diabetes: a systematic review. Cardiovascular Diabetology. 2022;21(1):272. doi: 10.1186/s12933-022-01703-5.

25. Hsu CC, Kanter JE, Kothari V, Bornfeldt KE. Quartet of APOCs and the Different Roles They Play in Diabetes. Arterioscler Thromb Vasc Biol. 2023;43(7):1124–33. Epub 20230525. doi: 10.1161/atvbaha.122.318290. PubMed PMID: 37226733; PMCID: PMC10330679.

26. Ellis D, Watanabe K, Wilmanski T, Lustgarten MS, Korat AVA, Glusman G, Hadlock JJ, Fiehn O, Sebastiani P, Price ND, Hood L, Magis AT, Evans SJ, Pflieger L, Lovejoy JC, Gibbons SM, Funk CC, Baloni P, Rappaport N. APOE Genotype and Biological Age Impact Inter-Omic Associations Related to Bioenergetics. bioRxiv. 2024. Epub 20241112. doi: 10.1101/2024.10.17.618322. PubMed PMID: 39605362; PMCID: PMC11601402.

27. Thambisetty M. Plasma Apolipoprotein E Levels and Risk of Dementia—You Are the Company You Keep. JAMA Network Open. 2020;3(7):e209501-e. doi: 10.1001/jamanetworkopen.2020.9501.

28. Li Z, Shue F, Zhao N, Shinohara M, Bu G. APOE2: protective mechanism and therapeutic implications for Alzheimer’s disease. Mol Neurodegener. 2020;15(1):63. Epub 20201104. doi: 10.1186/s13024-020-00413-4. PubMed PMID: 33148290; PMCID: PMC7640652.

29. Lumsden AL, Mulugeta A, Zhou A, Hypponen E. Apolipoprotein E (APOE) genotype-associated disease risks: a phenome-wide, registry-based, case-control study utilising the UK Biobank. EBioMedicine. 2020;59:102954. Epub 20200817. doi: 10.1016/j.ebiom.2020.102954. PubMed PMID: 32818802; PMCID: PMC7452404.

30. Kim H, Devanand DP, Carlson S, Goldberg TE. Apolipoprotein E Genotype e2: Neuroprotection and Its Limits. Front Aging Neurosci. 2022;14:919712. Epub 20220714. doi: 10.3389/fnagi.2022.919712. PubMed PMID: 35912085; PMCID: PMC9329577.

31. Döring Y, Drechsler M, Wantha S, Kemmerich K, Lievens D, Vijayan S, Gallo RL, Weber C, Soehnlein O. Lack of Neutrophil-Derived CRAMP Reduces Atherosclerosis in Mice. Circulation Research. 2012;110(8):1052–6. doi: doi:10.1161/CIRCRESAHA.112.265868.

32. N. SJ, H. SL. Myeloperoxidase and Cardiovascular Disease. Arteriosclerosis, Thrombosis, and Vascular Biology. 2005;25(6):1102–11. doi: doi:10.1161/01.ATV.0000163262.83456.6d.

33. Smyth LCD, Murray HC, Hill M, van Leeuwen E, Highet B, Magon NJ, Osanlouy M, Mathiesen SN, Mockett B, Singh-Bains MK, Morris VK, Clarkson AN, Curtis MA, Abraham WC, Hughes SM, Faull RLM, Kettle AJ, Dragunow M, Hampton B. Neutrophil-vascular interactions drive myeloperoxidase accumulation in the brain in Alzheimer’s disease. Acta Neuropathologica Communications. 2022;10(1):38. doi: 10.1186/s40478-022-01347-2.

34. Keragala CB, Medcalf RL. Plasminogen: an enigmatic zymogen. Blood. 2021;137(21):2881–9. doi: 10.1182/blood.2020008951.

35. Tavares LP, Negreiros-Lima GL, Lima KM, E Silva PMR, Pinho V, Teixeira MM, Sousa LP. Blame the signaling: Role of cAMP for the resolution of inflammation. Pharmacological Research. 2020;159:105030. doi: 10.1016/j.phrs.2020.105030.

36. Vandenberghe-Dürr S, Gilliet M, Di Domizio J. OLFM4 regulates the antimicrobial and DNA binding activity of neutrophil cationic proteins. Cell Reports. 2024;43(10). doi: 10.1016/j.celrep.2024.114863.

37. Brandt J, Spencer, M, & Folstein, M. The telephone interview for cognitive status.1988. 111–7 p.

38. Wojczynski MK, Jiuan Lin S, Sebastiani P, Perls TT, Lee J, Kulminski A, Newman A, Zmuda JM, Christensen K, Province MA. NIA Long Life Family Study: Objectives, Design, and Heritability of Cross-Sectional and Longitudinal Phenotypes. J Gerontol A Biol Sci Med Sci. 2022;77(4):717–27. doi: 10.1093/gerona/glab333. PubMed PMID: 34739053; PMCID: PMC8974329.

39. Reed ER, Chandler KB, Lopez P, Costello CE, Andersen SL, Perls TT, Li M, Bae H, Soerensen M, Monti S, Sebastiani P. Cross-platform proteomics signatures of extreme old age. Geroscience. 2024. Epub 20240725. doi: 10.1007/s11357-024-01286-x. PubMed PMID: 39048883.

40. Cox J, Mann M. MaxQuant enables high peptide identification rates, individualized p.p.b.-range mass accuracies and proteome-wide protein quantification. Nat Biotechnol. 2008;26(12):1367–72. doi: 10.1038/nbt.1511. PubMed PMID: 19029910.

41. Muller C, Schillert A, Rothemeier C, Tregouet DA, Proust C, Binder H, Pfeiffer N, Beutel M, Lackner KJ, Schnabel RB, Tiret L, Wild PS, Blankenberg S, Zeller T, Ziegler A. Removing Batch Effects from Longitudinal Gene Expression - Quantile Normalization Plus ComBat as Best Approach for Microarray Transcriptome Data. PLoS One. 2016;11(6):e0156594. doi: 10.1371/journal.pone.0156594. PubMed PMID: 27272489; PMCID: PMC4896498.

42. Benjamini Y, Hochberg Y. Controlling the false discovery rate - a practical and powerful approach to multiple testing. J Roy Statist Soc Series B 1995;57(1):289–300. PubMed PMID: ISI:A1995QE45300017.

43. Acharya S, Liao S, Jung WJ, Kang YS, Moghaddam VA, Feitosa MF, Wojczynski MK, Lin S, Anema JA, Schwander K, Connell JO, Province MA, Brent MR. A methodology for gene level omics-WAS integration identifies genes influencing traits associated with cardiovascular risks: the Long Life Family Study. Human Genetics. 2024;143(9):1241–52. doi: 10.1007/s00439-024-02701-1.

